# From variants to networks: resolving low SNP overlap in dairy cattle meta-analysis through multi-level functional mapping

**DOI:** 10.64898/2026.10.09.757896

**Authors:** M. Jakimowicz, D. Grabolus, T. Suchocki

## Abstract

The objective of this study was to perform a systematic review and multi-level meta-analysis of genome-wide association studies (GWAS) for functional traits in dairy cattle, focusing on calving performance and udder health. We hypothesized that despite the limited replication of individual SNPs across independent studies, functionally related variants would converge on common genes, biological pathways, and genomic regions. Following a systematic literature search and a two-stage filtering process, we extracted data from 20 independent articles covering seven traits: direct and maternal calving ease, direct and maternal stillbirth, clinical mastitis, somatic cell count, and somatic cell score. We first conducted a conventional SNP-level meta-analysis, which demonstrated minimal variant overlap across studies, preventing the detection of significant pooled effects. To resolve this low overlap, we mapped the associated variants to higher organizational levels, including genes, proteins, KEGG and Reactome pathways, and quantitative trait loci (QTL), and combined the statistical evidence using Fisher’s method. This multi-level functional mapping revealed significant biological convergence. At the gene level, we identified shared genes such as CCDC170 linking maternal calving ease and maternal stillbirth, while DCK, TMPRSS11F, and SLC4A4 were significant for udder health traits. Pathway-level meta-analysis highlighted major functional networks, demonstrating that mastitis-related traits were strongly associated with immune response, signal transduction, and cellular stress pathways, whereas maternal calving traits were dominated by endocrine and calcium signaling. Furthermore, our cross-trait overlap analysis identified 35 shared pathways and 30 shared QTL, reflecting a common biological architecture connecting calving difficulties and mastitis susceptibility. Notably, the innate immune system and signal transduction pathways consistently overlapped across multiple functional traits. In conclusion, conventional SNP-level meta-analysis has limited utility for integrating dairy cattle GWAS due to insufficient cross-study marker overlap. Moving from individual variants toward higher-order functional units effectively reveals reproducible genetic signals and functional convergence. The identified genes, biological pathways, and genomic regions offer valuable insights that can be leveraged as biological priors to improve genomic prediction models for dairy cattle health and reproduction.

**Interpretive summary:** Genetic studies on dairy cattle health and calving traits often show limited replication of individual variants across independent populations. This study addresses this limitation by moving beyond conventional single-marker meta-analysis and integrating genome-wide association data at higher levels of biological organization: genes, proteins, pathways, and quantitative trait loci. By mapping variants from 20 independent studies to these higher-order functional units, we revealed a shared biological architecture*-*such as immune and calcium signaling*-*linking calving difficulties with mastitis susceptibility. These findings provide biological priors that can enhance genomic prediction models and breeding programs for dairy cattle functional traits.

## INTRODUCTION

Functional traits such as calving performance, stillbirth, and udder health are important components of dairy cattle breeding because they affect animal health, reproductive performance, longevity, and overall production efficiency. These traits are typically complex and polygenic, with their phenotypic variation influenced by many genetic variants of small to moderate effect. Genome-wide association studies (GWAS) have provided an effective framework for identifying genomic regions associated with such complex traits; however, results obtained across studies are often difficult to compare directly because different populations, marker sets, sample sizes, and analytical approaches may lead to limited replication of individual SNP associations.

Meta-analysis offers a means of integrating evidence across independent GWAS and increasing the ability to identify reproducible genetic signals. Methods based on combined effect estimates or statistical evidence have therefore become an important component of the analysis of genome-wide association data (Evangelou and Ioannidis, 2013). Software such as METAL allows results from independent association studies to be combined using effect estimates and their corresponding standard errors (Willer *et al*., 2010). Nevertheless, conventional SNP-level meta-analysis relies on the same genetic variants being reported across studies. When overlap between reported SNPs is limited, biologically related signals located within the same genes, pathways, or quantitative trait loci (QTL) may remain undetected because they are represented by different variants in individual studies.

An alternative strategy is to move beyond individual variants and integrate GWAS evidence at higher levels of biological organization. Single nucleotide polymorphisms can be functionally linked to genes and their protein products and subsequently assigned to biological pathways and previously characterized QTL. Resources such as the Bovine Genome Database/BovineMine provide integrated information for bovine genomic annotation (Kambal *et al*., 2026), whereas KEGG and Reactome enable the interpretation of genomic findings within biological pathways and molecular processes (Kanehisa *et al*., 2025; Ragueneau *et al*., 2026). Such multi-level mapping may reveal biological convergence between studies even when the underlying associated variants are different. It may also facilitate identification of biological mechanisms shared across related traits.

Therefore, the objective of this study was to perform a systematic review and multi-level meta-analysis of GWAS results for functional traits in dairy cattle, including calving ease, stillbirth, clinical mastitis, mastitis, somatic cell count (SCC), and somatic cell score (SCS). We first evaluated the degree of overlap among reported SNP associations and performed conventional SNP-level meta-analysis where common variants were available. We then mapped associated variants to genes, proteins, KEGG and Reactome pathways, and QTL and combined statistical evidence at each level of biological organization. Finally, we evaluated cross-trait overlap to determine whether moving from individual variants toward higher-order functional units could reveal shared biological signals that were not detectable at the SNP level. We hypothesized that, despite limited replication of individual SNPs among GWAS, functionally related variants would converge on common genes, pathways, and genomic regions, thereby revealing a shared biological architecture of dairy cattle functional traits.

## MATERIALS AND METHODS

### Study Selection

We conducted a systematic review and an aggregate data meta-analysis of SNPs obtained from GWAS for functional traits in dairy cattle. We focused our search on the following traits: calving ease (CE), direct calving ease (DCE), maternal calving ease (MCE), stillbirth (SB), direct stillbirth (DSB), maternal stillbirth (MSB), clinical mastitis (CM), mastitis, SCS, and SCC. We searched PubMed (https://pubmed.ncbi.nlm.nih.gov/) and Web of Science (WoS; https://www.webofscience.com) between April 2025 and February 2026. Two authors independently searched the databases: DG screened PubMed, MJ screened WoS, and TS resolved any discrepancies. We included only full-text articles available for retrieval. Search queries used to retrieve studies are presented in Supplemental File S1.

### Article Filtering

After combining the results from both databases, we filtered the data using a two-stage process: first at the SNP level, and second at the article level. At the SNP level, we excluded variants that: 1) lacked a *P*-value in numerical form (e.g., reported as *P < 0.001*); 2) lacked a reported effect size; 3) lacked a unique SNP identifier; or 4) had an effect marked only as chromosome-wide. After removing variants based on these criteria, we proceeded to article-level filtering. Based on consensus of the working group, we excluded articles that met any of the following criteria: 1) duplicate publications; 2) lack of reported effect size for all analyzed SNPs; 3) sample size fewer than 150 animals; or 4) lack of RefSNP (rs) accession numbers for all SNPs. We established a threshold of 150 individuals to balance study retention and data reliability. Because the included studies spanned a long period (2008 to 2024), sample sizes varied substantially over time. Furthermore, traits such as CM are less frequently analyzed in the literature; excluding smaller studies would have unnecessarily restricted the dataset rather than improving analytical rigor. Additionally, we excluded an entire trait from meta-analysis if fewer than 3 studies remained after filtering. All filtering stages were performed using R version 4.5.2 (R Core Team, 2025).

### Statistical Analysis

First, we mapped SNP names lacking rs numbers to their corresponding rs identifiers based on the ARS-UCD2.0 reference genome using BovineMine v1.7 (Kambal *et al*., 2026). After unifying all SNP identifiers, we created a separate dataset for each trait and transformed extracted *P*-values into their raw numerical form. Trait-specific meta-analyses were then conducted using METAL software (Willer *et al*., 2010), applying a fixed-effects model with inverse-variance weighting based on SNP effect sizes and corresponding standard errors.

### Functional Annotation and Higher-Level Meta-Analysis

Following the initial mollows a chi-square distribution mapped to genes, proteins, biological pathways (KEGG; Kanehisa *et al*., 2025; Reactome; Ragueneau *et al*., 2026), and QTL using BovineMine v1.7.Separate meta-analyses were conducted for each higher organizational level. Because source articles lacked consistent effect size measures for identical functional units and the number of studies per unit was small, we applied Fisher’s method to pool independent *P*-values across studies (Evangelou and Ioannidis, 2013). The test statistic was defined as:

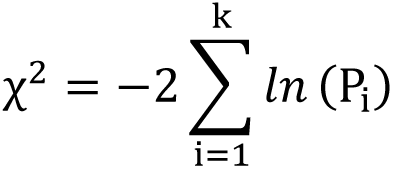

where *P_i_* represents the *P*-value for the *i*-th study, and *k* is the number of studies in the meta-analysis. Under the null hypothesis, *χ*^2^ follows a chi-square distribution with 2*k* degrees of freedom. To satisfy the assumption of independence required by Fisher’s method, we selected a single representative *P*-value specifically the lowest *P*-value per study for each biological entity. This approach allowed us to consolidate data and identify the strongest association signal at each organizational level. All higher-level meta-analyses were performed using custom R scripts.

## RESULTS

The initial database search yielded 61 potential articles after title and abstract screening. Full-text retrieval and multi-stage filtering narrowed this selection to 20 articles for final analysis. The identification process was conducted according to PRISMA 2020 guidelines (Page *et al*., 2021), as illustrated in Figure 1. The included papers were published between 2008 and 2024, capturing over 15 years of dairy cattle genomic research. Table 1 summarizes article counts before and after filtration by trait. Direct stillbirth was the only trait without article loss after filtering. In contrast, CE and SB were excluded due to lack of eligible studies, and mastitis was excluded due to an insufficient study count (*n = 2*). Subsequent analyses focused on 7 traits. Table 2 presents the characteristics of the included studies. Population sizes varied significantly, ranging from 350 to 294,079 animals. Similarly, the number of analyzed SNPs per study ranged from 34 to 966. Full study characteristics are provided in Supplemental Table S2.

**Figure 1.**
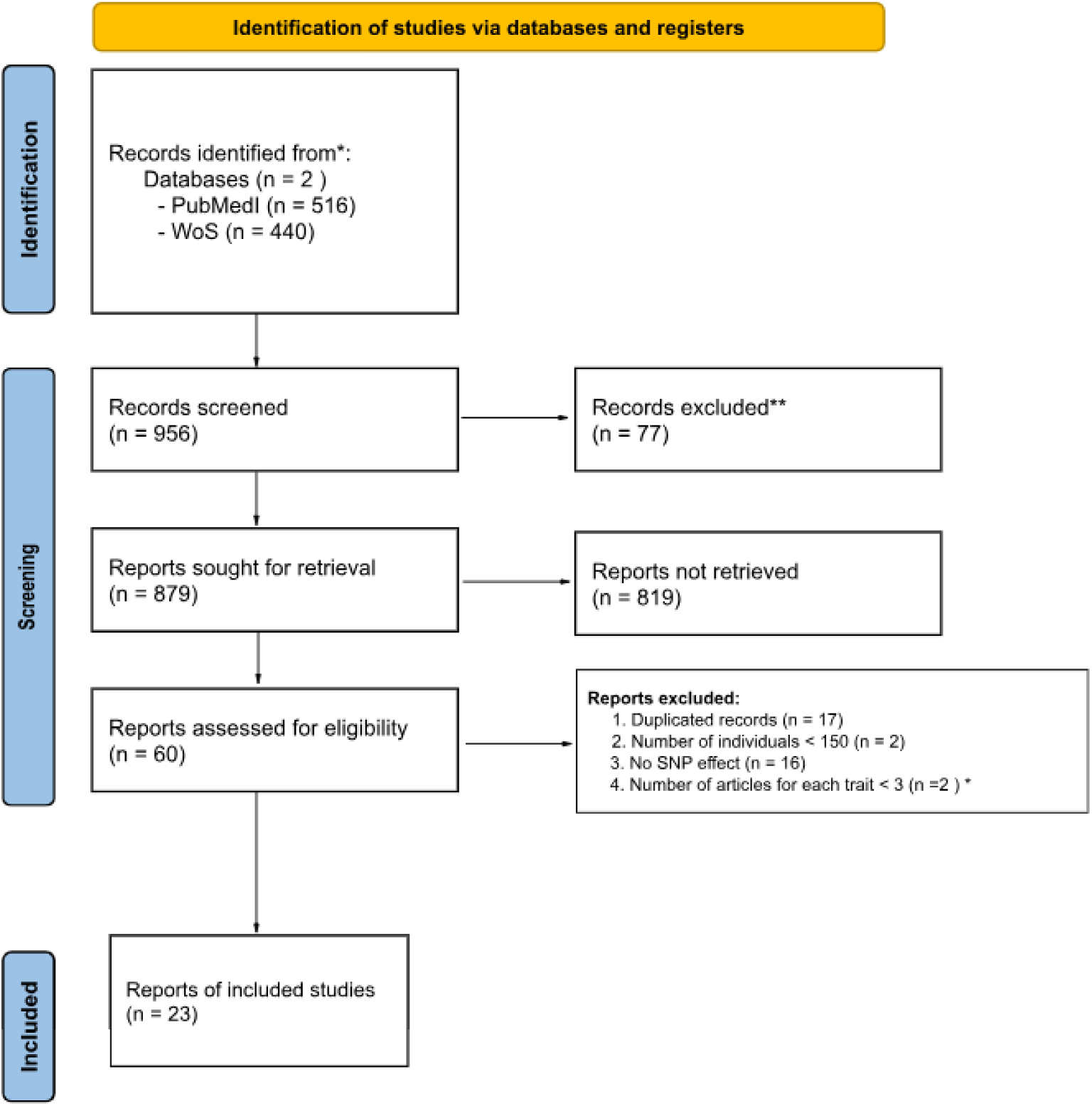
PRISMA 2020 flow diagram illustrating the identification, screening, and selection process for genome-wide association studies included in the multi-level meta-analysis. * This step aimed to exclude entire traits that failed to meet the threshold. Consequently, we removed two articles (n=2), resulting in the elimination of two traits.

**Table 1.** Article counts before and after filtration by trait.

| Trait | No. of articles before filtration | No. of articles after filtration |
| --- | --- | --- |
| CE | 3 | 0 |
| DCE | 6 | 4 |
| DSB | 4 | 4 |
| MCE | 5 | 3 |
| MSB | 3 | 3 |
| SB | 4 | 0 |
| SCC | 7 | 3 |
| SCS | 16 | 5 |
| clinical mastitis | 9 | 6 |
| mastitis | 7 | 2 |

**Table 2.** Characteristics of the studies included in the meta-analysis.

| DOI | Traits | Number of animals | Number of SNPs for each trait |
| --- | --- | --- | --- |
| https://doi.org/10.1038/s41598-020-64575-3 | SCS | 1,152 | 182 |
| https://doi.org/10.1111/age.12053 | Clinical mastitis | 2,098 | 80 |
| https://doi.org/10.1111/age.12624 | Clinical mastitis | 1,741 | 80 |
| https://doi.org/10.1111/j.1439-0388.2008.00793.x | SCS | 462 | 182 |
| https://doi.org/10.1186/1471-2156-13-21 | Clinical mastitis | 914 | 80 |
| https://doi.org/10.1186/1471-2156-14-49 | SCS | 550 | 182 |
| https://doi.org/10.1186/s12864-024-10782-5 | DCE | 252,285 | 100 |
| https://doi.org/10.1186/s12864-024-10782-5 | MCE | 252,285 | 297 |
| https://doi.org/10.1186/s12864-024-10782-5 | DSB | 252,285 | 49 |
| https://doi.org/10.1186/s12864-024-10782-5 | MSB | 252,285 | 966 |
| https://doi.org/10.1371/journal.pone.0050137 | DCE | 323 | 100 |
| https://doi.org/10.1534/genetics.110.124057 | DCE | 1,800 | 100 |
| https://doi.org/10.1534/genetics.110.124057 | DSB | 1,800 | 49 |
| https://doi.org/10.3168/jds.2010-3381 | DSB | 2,531 | 49 |
| https://doi.org/10.3168/jds.2013-7149 | SCS | 1,412 | 182 |
| https://doi.org/10.3168/jds.2014-8141 | Clinical mastitis | 4,442 | 6 |
| https://doi.org/10.3168/jds.2014-8141 | Clinical mastitis | 1,126 | 6 |
| https://doi.org/10.3168/jds.2014-8141 | Clinical mastitis | 4,992 | 6 |
| https://doi.org/10.3168/jds.2014-8141 | SCC | 4,992 | 6 |
| https://doi.org/10.3168/jds.2017-14067 | Clinical mastitis | 32,491 | 80 |
| https://doi.org/10.3168/jds.2020-18209 | Clinical mastitis | 1,062 | 80 |
| https://doi.org/10.3168/jds.2024-25978 | MCE | 32,352 | 297 |
| https://doi.org/10.3168/jds.2024-25978 | MSB | 32,352 | 966 |
| https://doi.org/10.3389/fgene.2018.00141 | Mastitis | 993 | 34 |
| https://doi.org/10.3389/fgene.2018.00141 | SCC | 993 | 34 |
| https://doi.org/10.3389/fgene.2019.00412 | SCS | 294,079 | 182 |
| https://doi.org/10.3389/fvets.2022.932034 | SCS | 1,313 | 182 |
| https://doi.org/10.3390/ani12091127 | DCE | 30,603 | 100 |
| https://doi.org/10.3390/ani12091127 | MCE | 29,738 | 297 |
| https://doi.org/10.3390/ani12091127 | DSB | 24,521 | 49 |
| https://doi.org/10.3390/ani12091127 | MSB | 28,081 | 966 |
| https://doi.org/10.3390/ani14182713 | Mastitis | 350 | 34 |

### SNP-Level Meta-Analysis

The conventional SNP-level meta-analysis revealed minimal marker overlap across studies for all traits. Only MCE and MSB possessed variants reported in more than 1 publication; for both traits, overlapping variants occurred within the exact same pair of articles. Two shared SNPs were identified for MCE (rs43608567 and rs43610539), and 1 shared SNP was identified for MSB (rs43610539). Despite their presence in multiple studies, pooled effect sizes calculated by METAL did not achieve statistical significance.

### Functional Annotation and Higher-Level Meta-Analysis

Because the SNP-level approach demonstrated limited utility due to insufficient marker overlap, we integrated associations at higher organizational levels: genes, proteins, KEGG pathways, Reactome pathways, and QTL. This mapped the analyzed SNPs to a total of 328 genes, 1,068 proteins, 502 KEGG pathways, 881 Reactome pathways, and 3,609 QTL. The distribution of mapped biological entities across traits is presented in Table 3.

**Table 3.** Number of mapped biological entities by trait.

| Trait | Number of genes | Numbers of proteins | Number of KEGG pathway | Number of reactome pathways | Number of QTL |
| --- | --- | --- | --- | --- | --- |
| DCE | 51 | 146 | 85 | 86 | 735 |
| MCE | 37 | 130 | 93 | 161 | 216 |
| DSB | 21 | 46 | 21 | 21 | 332 |
| MSB | 43 | 144 | 54 | 124 | 127 |
| clinical mastitis | 41 | 155 | 85 | 135 | 903 |
| SCC | 18 | 78 | 23 | 33 | 71 |
| SCS | 100 | 303 | 124 | 292 | 1166 |

### Gene Level

Gene-level meta-analysis yielded statistically significant results for 4 traits: MCE, MSB, SCC, and CM (Table 4). For MCE, 4 significant genes were identified: estrogen receptor 1 (ESR1), protocadherin-11 X-linked (LOC538674), coiled-coil domain containing 170 (CCDC170), and ENSBTAG00000001911 (a novel gene whose human ortholog is PCDH11X). For CM, 2 significant genes were identified: deoxycytidine kinase (DCK) and transmembrane serine protease 11F (TMPRSS11F). Single significant genes were identified for MSB (CCDC170) and SCC (SLC4A4).

**Table 4.** Genes significant in meta-analysis for each trait.

| Trait | Gene symbol | Effect | P-value |
| --- | --- | --- | --- |
| MCE | ESR1 | 3.536 | $1.491 \times 10^{-14}$ |
| | LOC538674 | 1.862 | $5.764 \times 10^{-14}$ |
| | CCDC170 | 2.935 | $7.472 \times 10^{-8}$ |
| | ENSBTAG00000001911 | 1.862 | $5.764 \times 10^{-14}$ |
| MSB | CCDC170 | 3.959 | $4.008 \times 10^{-14}$ |
| SCC | SLC4A4 | -3.015 | $2.201 \times 10^{-16}$ |
| CM | DCK | -0.500 | $3.318 \times 10^{-10}$ |
| | TMPRSS11F | -0.363 | $1.180 \times 10^{-8}$ |

### Protein Level

At the protein level, meta-analysis yielded statistically significant associations across 6 traits: MCE, SCC, CM, DSB, MSB, and SCS (Table 5). For MCE, 4 significant proteins were identified: estrogen receptor, estrogen receptor 1, cadherin domain-containing protein, and coiled-coil domain containing 170. For SCC, anion exchange protein and electrogenic sodium bicarbonate cotransporter 1 were significant. For CM, transmembrane protease serine and deoxycytidine kinase were significant. Single significant proteins were identified for DSB (KRAB domain-containing protein), MSB (coiled-coil domain containing 170), and SCS (RNA-directed DNA polymerase).

**Table 5.**
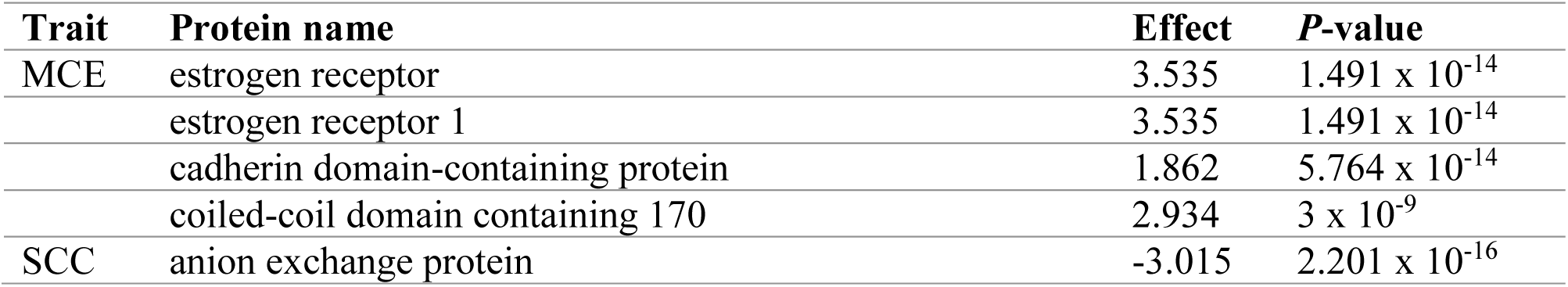
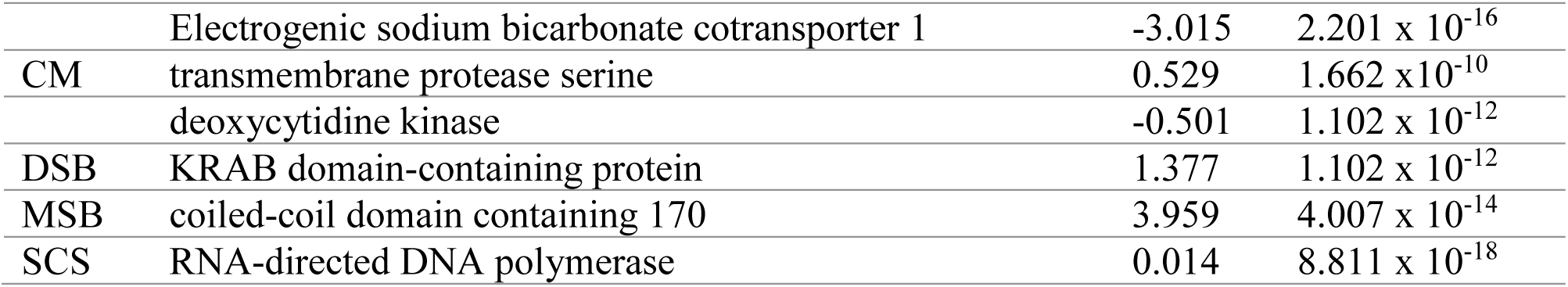
Proteins significant in meta-analysis for each trait.

### Pathway Level

KEGG Pathways. For CM, 32 KEGG pathways were significantly enriched (*P* < 0.05). Three pathways (bta00230, bta04360, bta04150) were identified across 3 independent publications, whereas 29 pathways appeared in 2 studies. Signaling pathways represented the largest functional category, accounting for 34% of all significant CM pathways. The mitogen-activated protein kinase (MAPK) signaling pathway (bta04010) contained the highest density of mapped genetic variants (10 SNPs; Figure 1).For SCS, 21 KEGG pathways were significant (Figure 2). Three pathways were reported across 3 studies (bta04727, bta04081, bta04150), and 18 pathways were reported in 2 studies. The neuroactive ligand-receptor interaction pathway (bta04080) captured the highest variant density (14 SNPs). Functional clustering highlighted processes directly related to immune response and leukocyte transport, including hematopoietic cell lineage (bta04640), chemokine signaling (bta04062), and cell adhesion molecules (bta04514). For MCE, 12 KEGG pathways were significant (Figure 3), with 3 pathways supported by 3 studies. Endocrine and calcium homeostasis dominated the functional profile, including estrogen signaling (bta04915), prolactin signaling (bta04917), and calcium signaling (bta04020).KEGG meta-analyses for DCE, DSB, MSB, and SCC revealed 4, 1, 3, and 4 significant pathways, respectively (Figure 4). For DCE, calcium signaling (bta04020) exhibited the highest variant density (8 SNPs). For DSB, spinocerebellar ataxia (bta05017) was the sole significant pathway (2 SNPs). For MSB, 3 pathways reached significance, led by neuroactive ligand-receptor interaction (bta04080). For SCC, 4 pathways were significant, with PI3K-Akt signaling (bta04151) capturing 14 SNPs. Reactome pathway meta-analysis for CM identified 31 significant pathways (Figure 5). Two pathways were present across 4 articles, 2 across 3 articles, and 27 across 2 articles. Signal transduction (R-BTA-162582) showed the highest variant density (52 SNPs). The pathways grouped into 6 primary functional categories: 1) immune system; 2) signal transduction and communication; 3) specific immune receptors and downstream cascades; 4) metabolism and nucleotide salvage; 5) transport and physiological systems; and 6) cellular stress responses. For MCE, 40 Reactome pathways were significant (Figure 6), with 8 pathways supported by 3 studies. Signal transduction (R-BTA-162582) contained 52 mapped SNPs across 3 articles. Functional groups included kinase signaling cascades, post-translational protein modification, estrogen and nuclear receptor signaling, gene expression regulation, innate/adaptive immunity, and cellular stress. For SCS, 37 Reactome pathways were significant (Figure 7), with 3 pathways present in 4 studies and 10 in 3 studies. Signal transduction (R-BTA-162582) represented a central biological node (52 SNPs across 4 studies). Primary functional clusters included master signal transduction, lipid metabolism, GPCR signaling, gene expression, and stress responses. Consolidated Reactome meta-analysis results for DCE, MSB, and SCC are shown in Figure 8; no Reactome pathway reached significance for DSB. For DCE, 3 pathways associated with immune system processes were significant (R-BTA-1280218, R-BTA-168256, R-BTA-168249). For MSB, 7 pathways were significant, led by signal transduction (R-BTA-162582; 52 SNPs). For SCC, 8 pathways were significant, clustering into transmembrane transport and immune response mechanisms.

**Figure 2.**
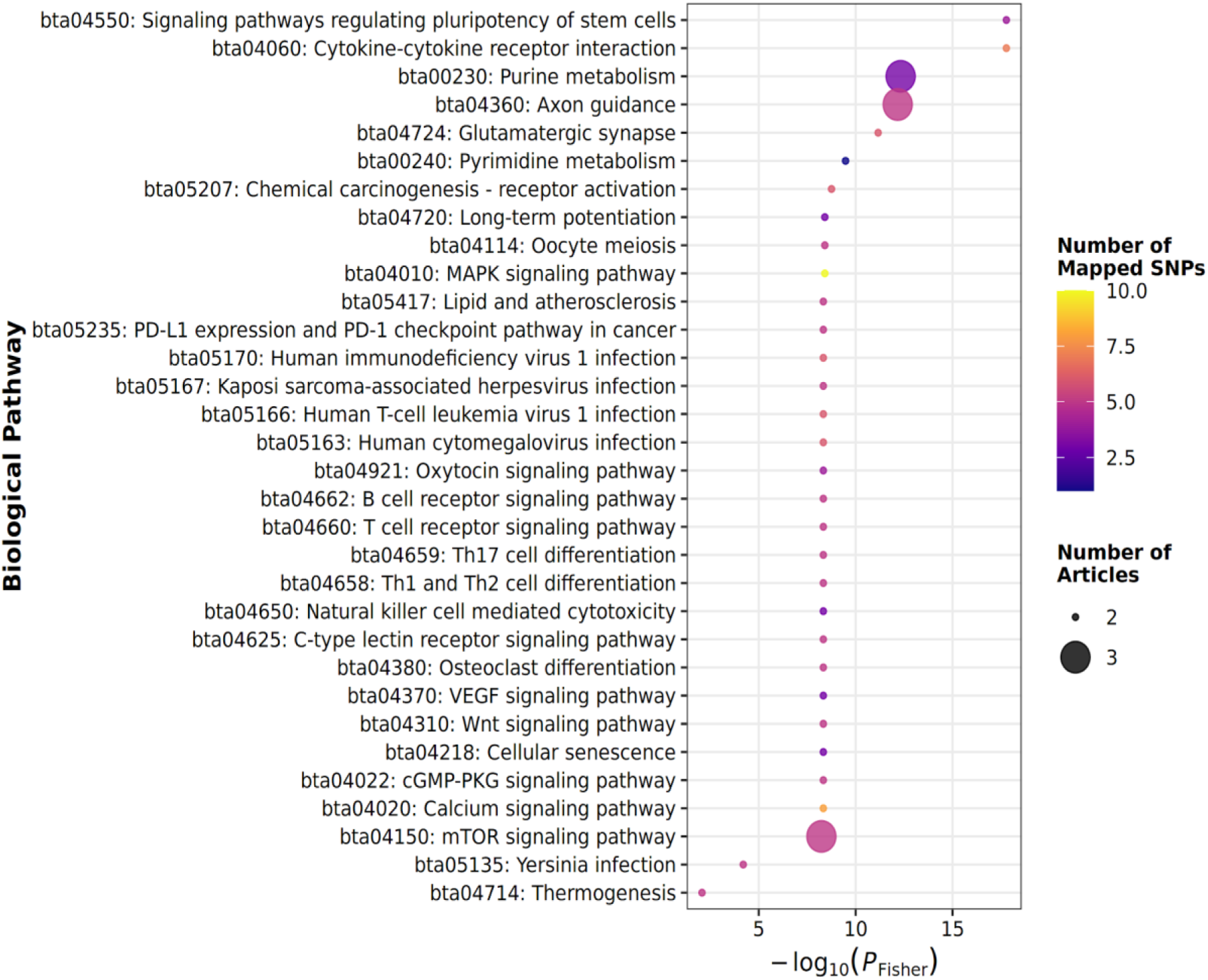
Bubble plot showing significant KEGG pathways identified in clinical mastitis meta-analysis. Bubble size indicates the number of contributing articles, and color intensity corresponds to the number of mapped single nucleotide polymorphisms (SNPs).

**Figure 3.**
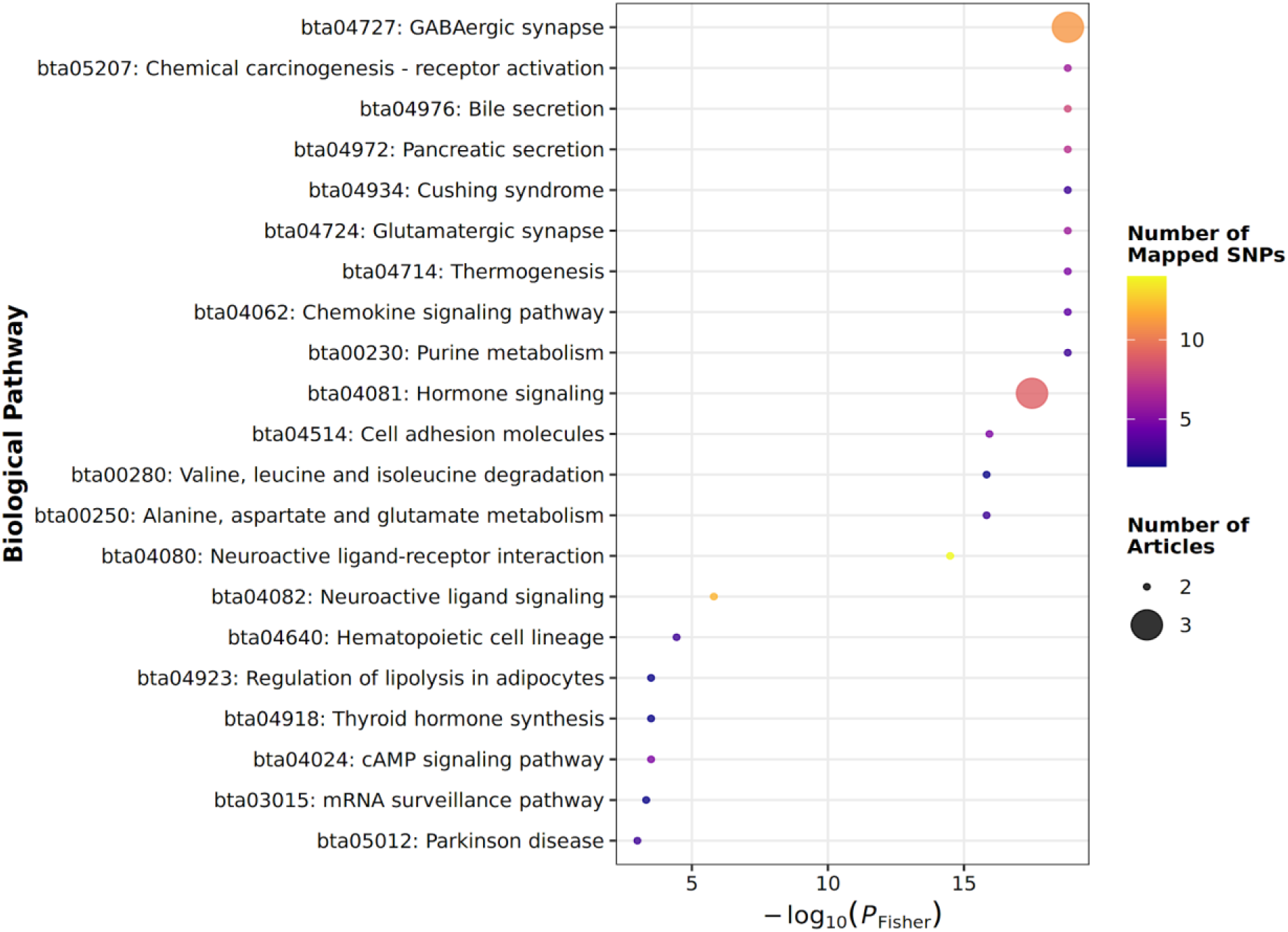
Bubble plot showing significant KEGG pathways identified in somatic cell score (SCS) meta-analysis.

**Figure 4.**
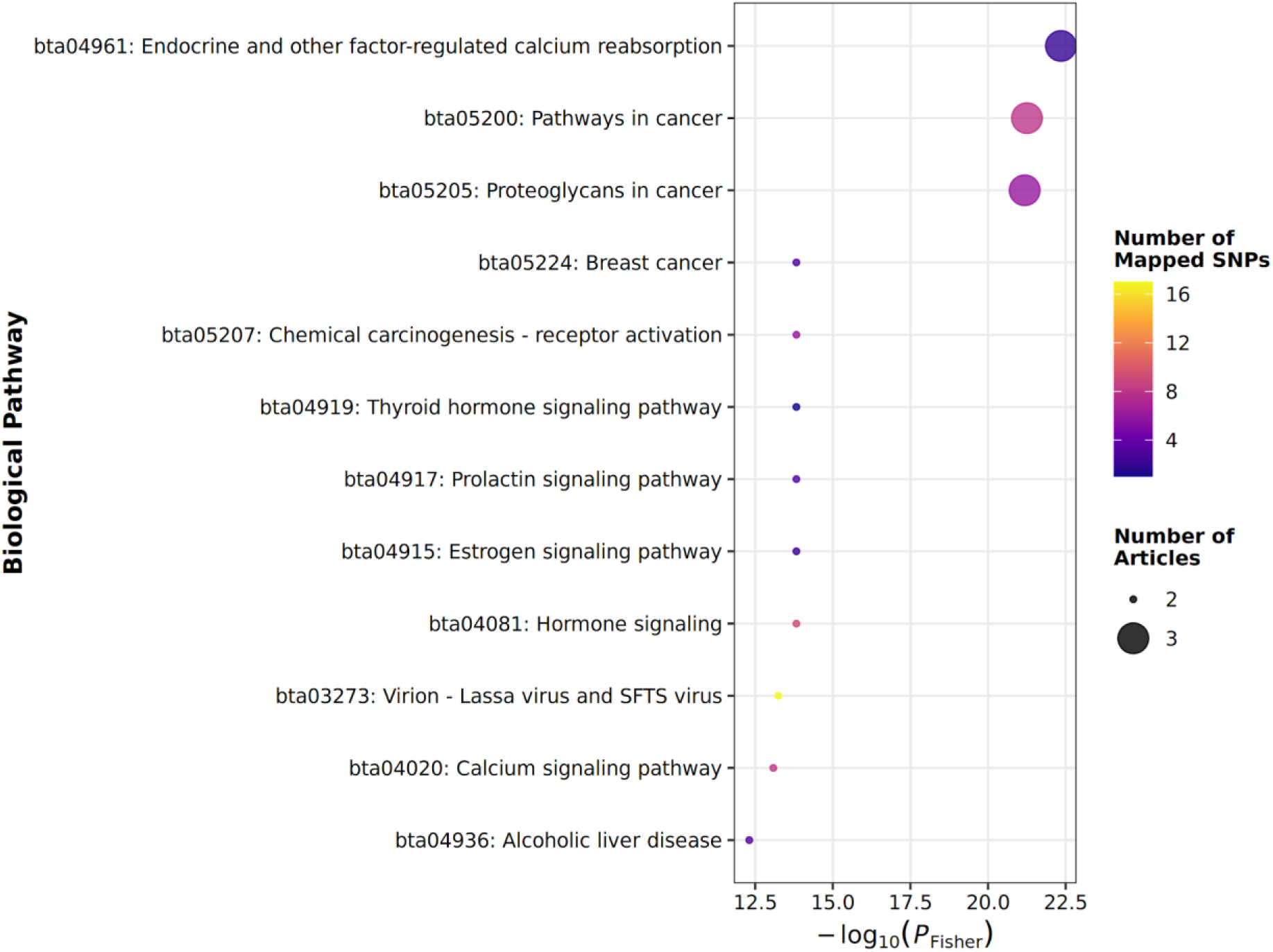
Bubble plot showing significant KEGG pathways identified in maternal calving ease (MCE) meta-analysis.

**Figure 5.**
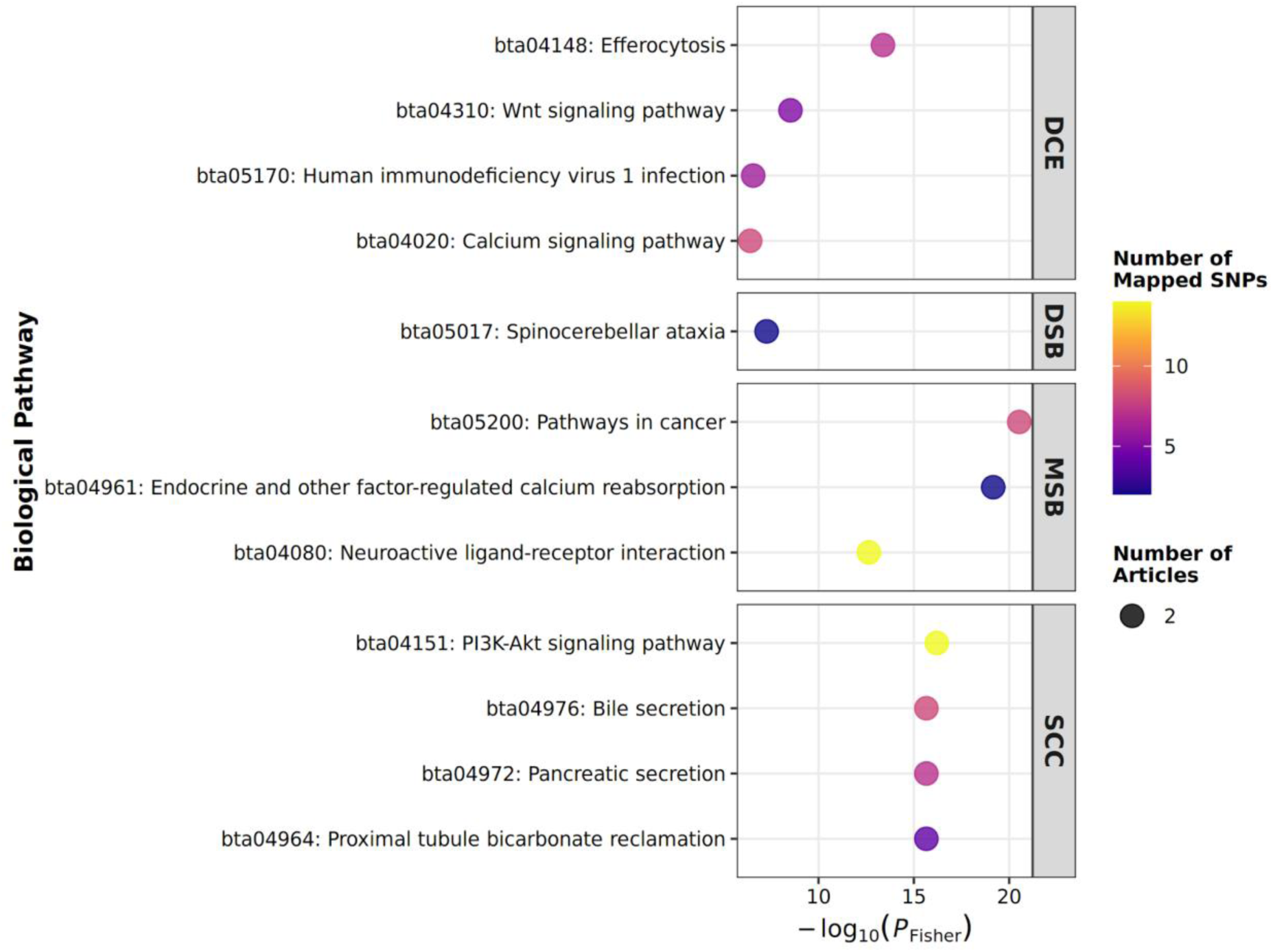
Combined bubble plot showing significant KEGG pathways across direct calving ease (DCE), direct stillbirth (DSB), maternal stillbirth (MSB), and somatic cell count (SCC) meta-analyses.

**Figure 6.**
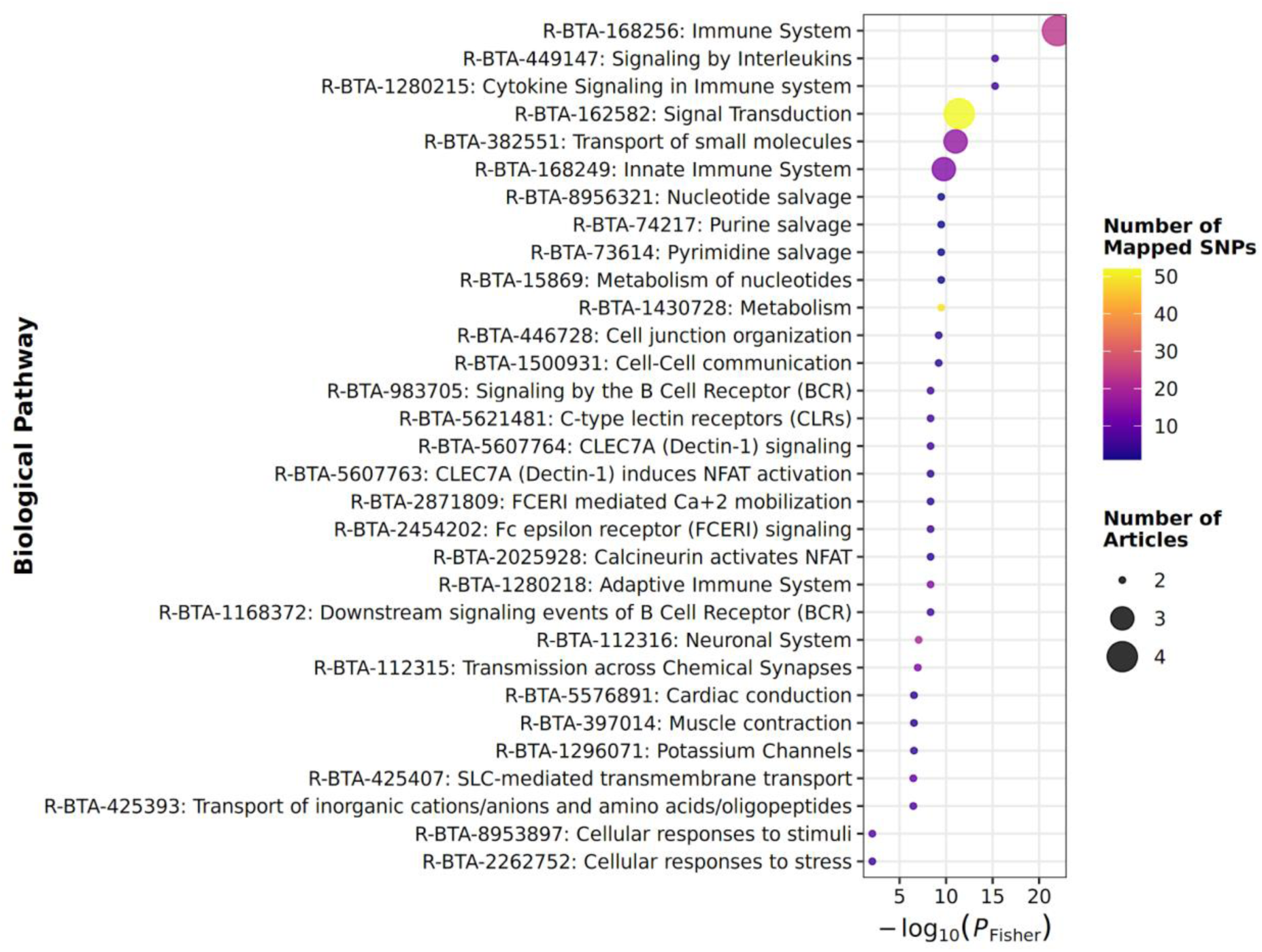
Bubble plot showing significant Reactome pathways identified in clinical mastitis meta-analysis.

**Figure 7.**
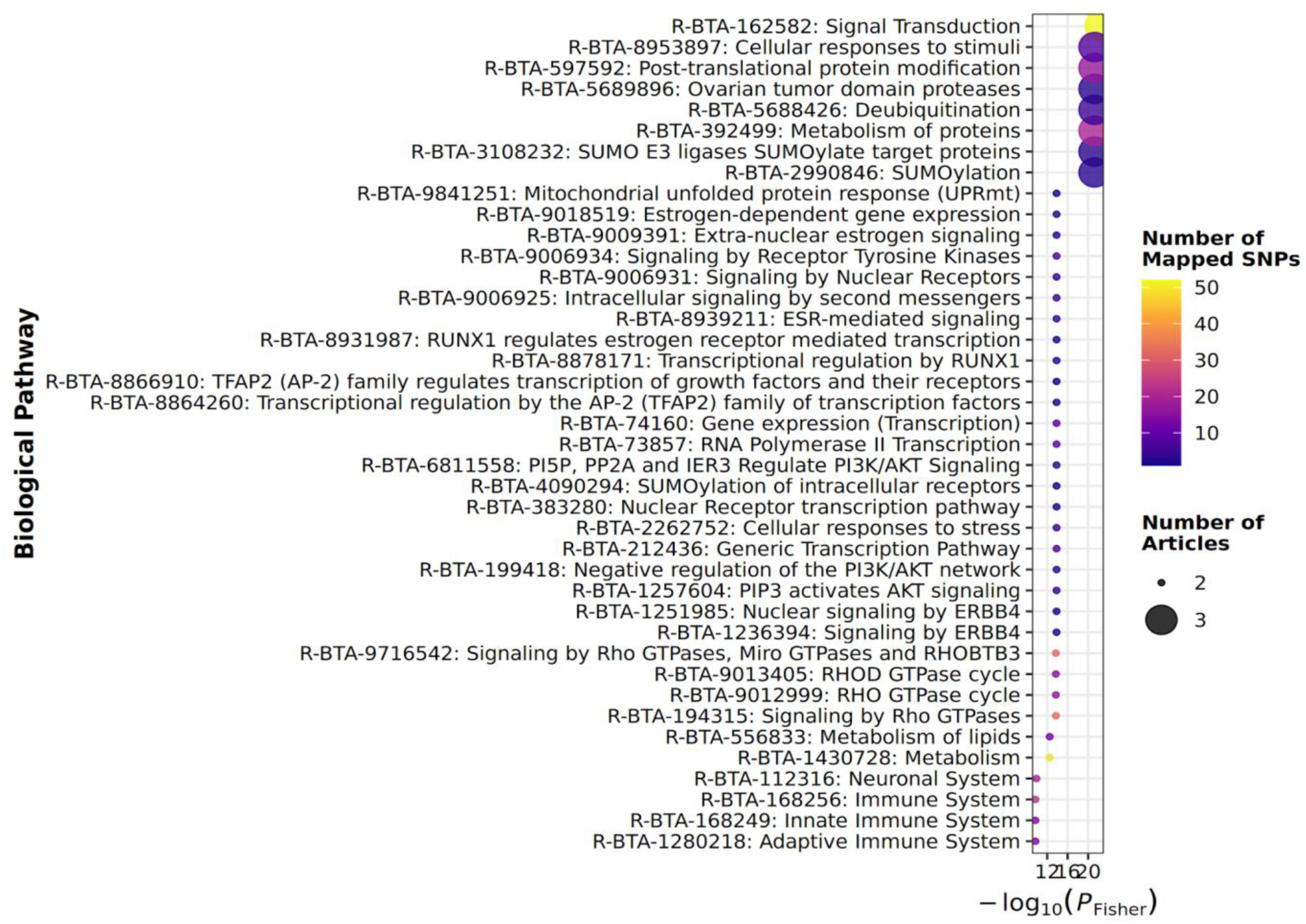
Bubble plot showing significant Reactome pathways identified in maternal calving ease (MCE) meta-analysis.

**Figure 8.**
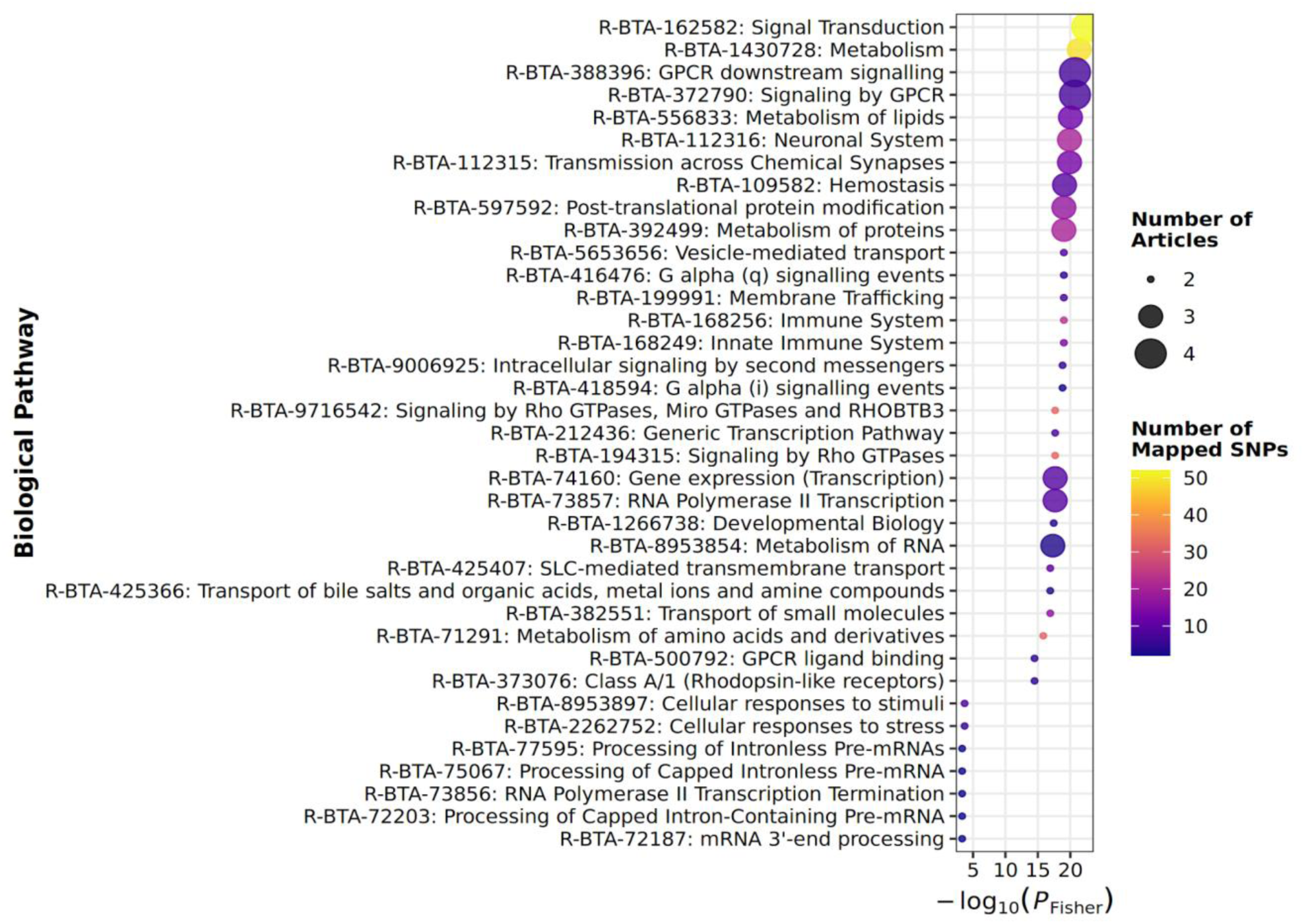
Bubble plot showing significant Reactome pathways identified in somatic cell score (SCS) meta-analysis.

### QTL Level

Meta-analysis at the QTL level identified 196 significant QTL across traits (Table 6), ranging from 4 for MSB to 85 for CM. Significant QTL were distributed across 11 chromosomes (Figure 9).

**Figure 9.**
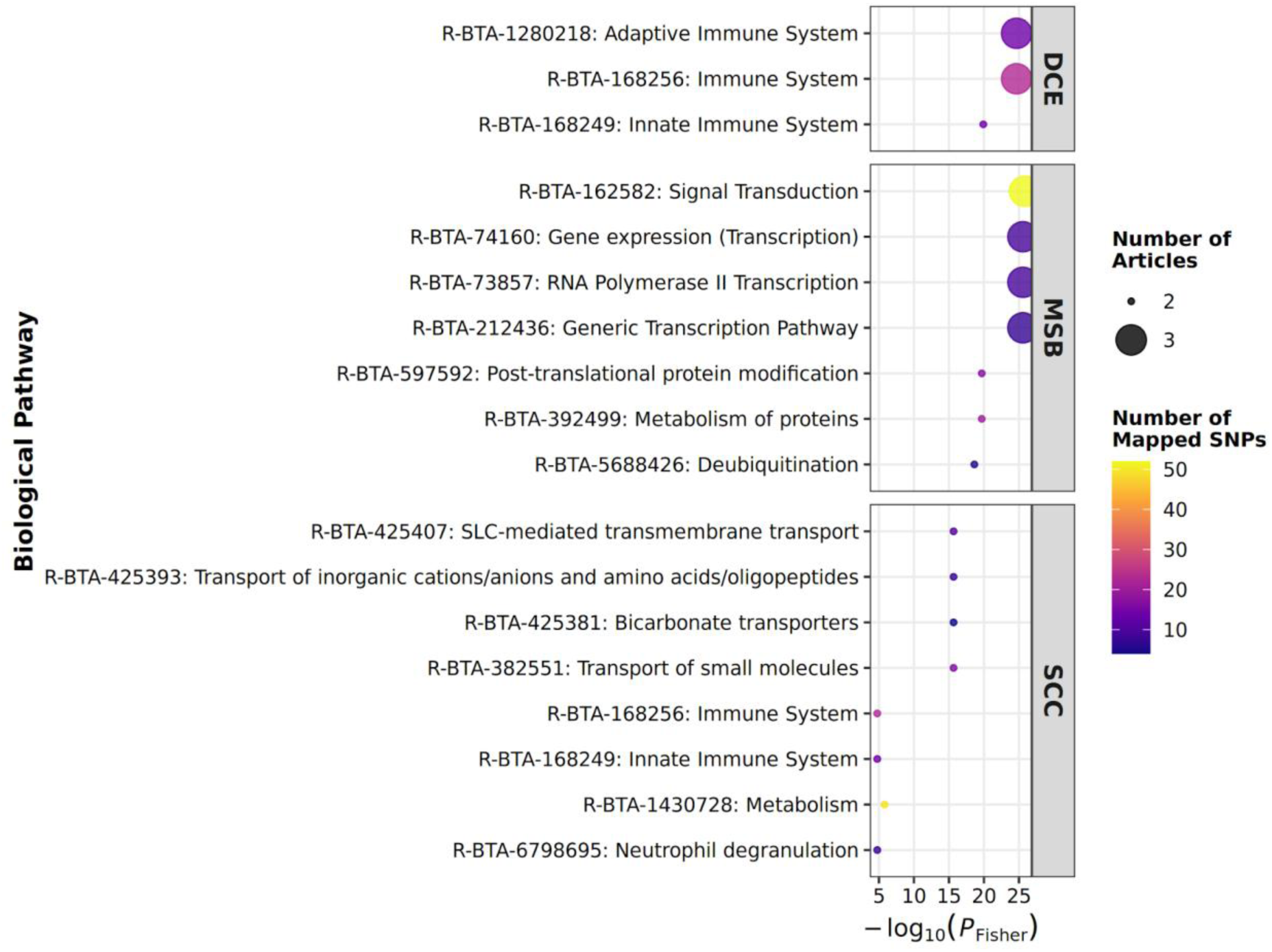
Combined bubble plot showing significant Reactome pathways across direct calving ease (DCE), maternal stillbirth (MSB), and somatic cell count (SCC) meta-analyses.

**Table 6.** Number of QTL significant for each trait.

| Trait | N QTL |
| --- | --- |
| clinical mastitis | 85 |
| DCE | 14 |
| DSB | 18 |
| SCS | 58 |
| SCC | 5 |
| MSB | 4 |
| MCE | 12 |

Loci clustered within 2 major genomic hotspots. The first hotspot on BTA6 (83.1 to 87.0 Mb) contained 56 QTL associated with CM, SCC, and MCE. The second hotspot on BTA14 (0.2 to 0.5 Mb) contained 47 QTL significant for SCS and milk production traits. Additionally, BTA18 harbored a prominent cluster of reproductive QTL (13.3 to 57.1 Mb). Remaining chromosomes exhibited localized overlaps affecting milk fat composition (BTA5, 19, 26) and health intervals (BTA16). A summary of significant QTL is provided in Table 7.

**Table 7.**
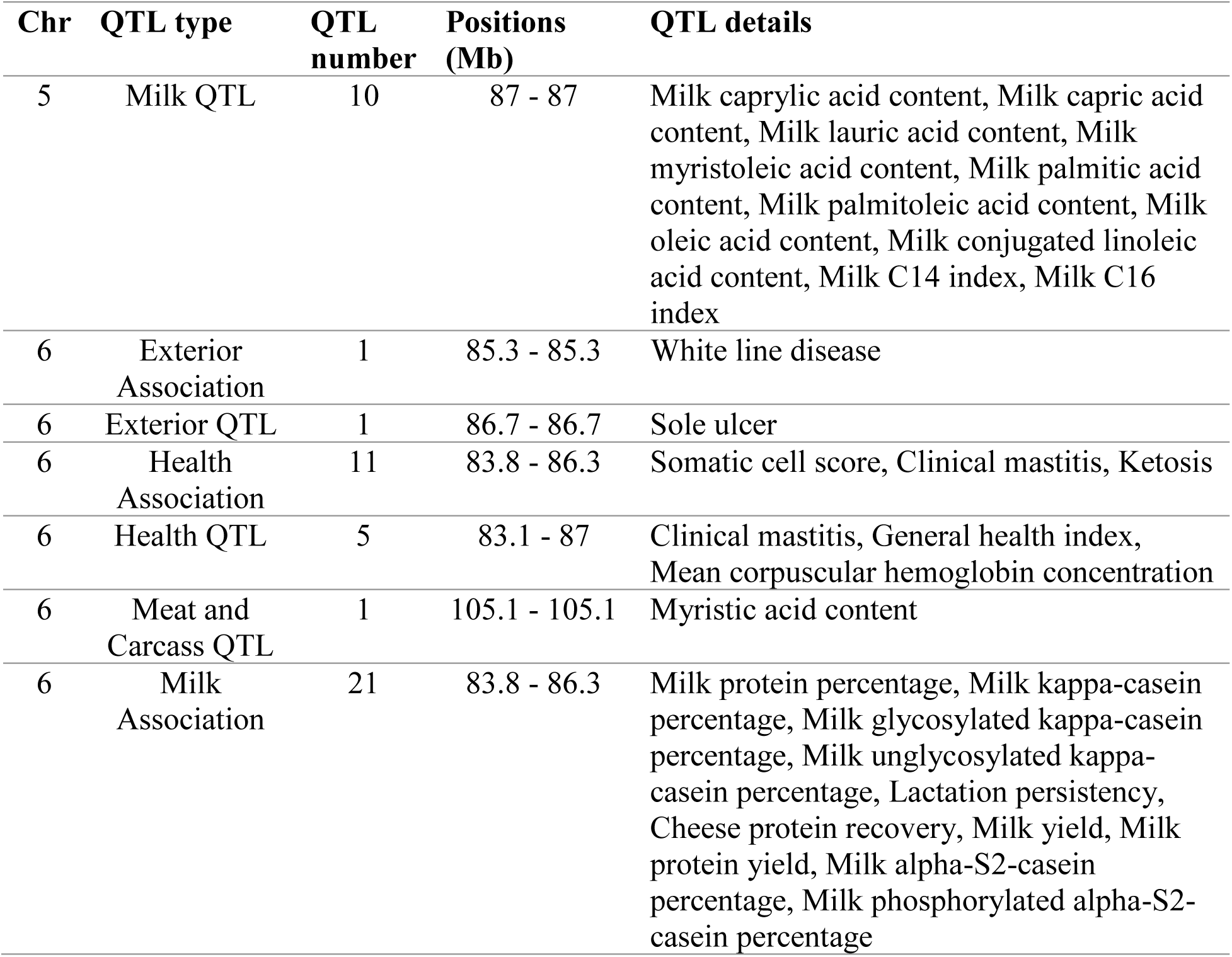
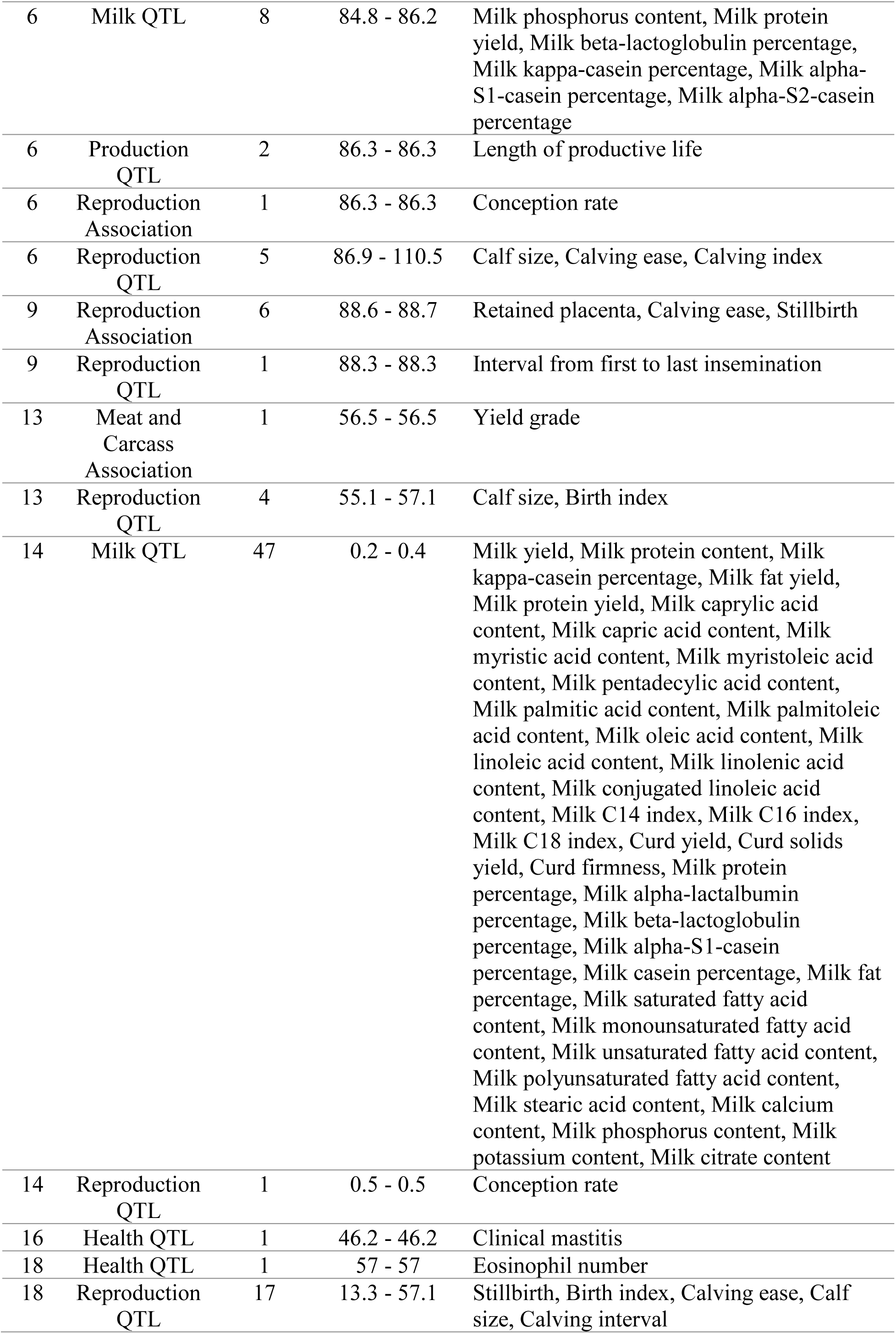
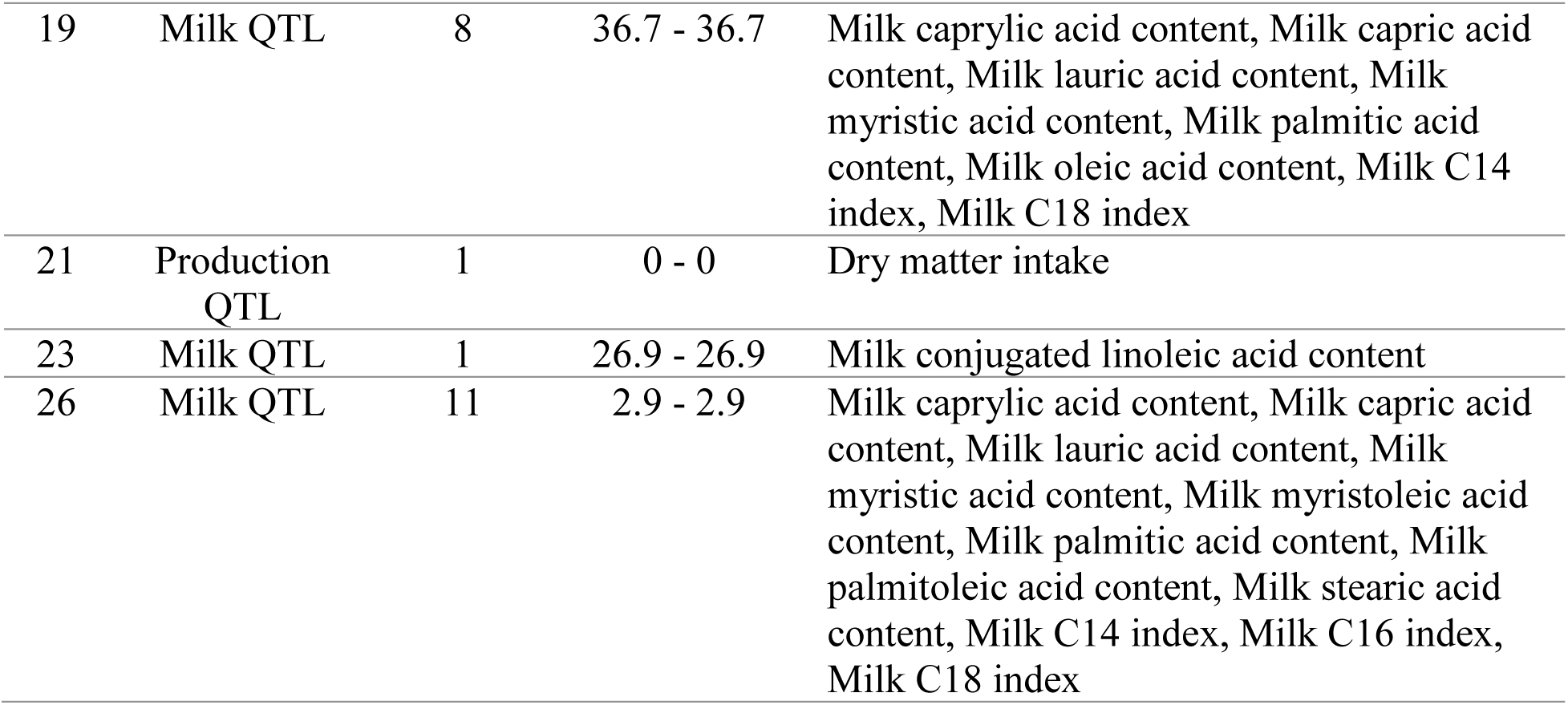
Summary of significant QTL for each trait.

| Chr | QTL type | QTL number | Positions (Mb) | QTL details |
| --- | --- | --- | --- | --- |
| 5 | Milk QTL | 10 | 87 - 87 | Milk caprylic acid content, Milk capric acid content, Milk lauric acid content, Milk myristoleic acid content, Milk palmitic acid content, Milk palmitoleic acid content, Milk oleic acid content, Milk conjugated linoleic acid content, Milk C14 index, Milk C16 index |
| 6 | Exterior Association | 1 | 85.3 - 85.3 | White line disease |
| 6 | Exterior QTL | 1 | 86.7 - 86.7 | Sole ulcer |
| 6 | Health Association | 11 | 83.8 - 86.3 | Somatic cell score, Clinical mastitis, Ketosis |
| 6 | Health QTL | 5 | 83.1 - 87 | Clinical mastitis, General health index, Mean corpuscular hemoglobin concentration |
| 6 | Meat and Carcass QTL | 1 | 105.1 - 105.1 | Myristic acid content |
| 6 | Milk Association | 21 | 83.8 - 86.3 | Milk protein percentage, Milk kappa-casein percentage, Milk glycosylated kappa-casein percentage, Milk unglycosylated kappa-casein percentage, Lactation persistency, Cheese protein recovery, Milk yield, Milk protein yield, Milk alpha-S2-casein percentage, Milk phosphorylated alpha-S2-casein percentage |
| 6 | Milk QTL | 8 | 84.8 - 86.2 | Milk phosphorus content, Milk protein yield, Milk beta-lactoglobulin percentage, Milk kappa-casein percentage, Milk alpha-S1-casein percentage, Milk alpha-S2-casein percentage |
| 6 | Production QTL | 2 | 86.3 - 86.3 | Length of productive life |
| 6 | Reproduction Association | 1 | 86.3 - 86.3 | Conception rate |
| 6 | Reproduction QTL | 5 | 86.9 - 110.5 | Calf size, Calving ease, Calving index |
| 9 | Reproduction Association | 6 | 88.6 - 88.7 | Retained placenta, Calving ease, Stillbirth |
| 9 | Reproduction QTL | 1 | 88.3 - 88.3 | Interval from first to last insemination |
| 13 | Meat and Carcass Association | 1 | 56.5 - 56.5 | Yield grade |
| 13 | Reproduction QTL | 4 | 55.1 - 57.1 | Calf size, Birth index |
| 14 | Milk QTL | 47 | 0.2 - 0.4 | Milk yield, Milk protein content, Milk kappa-casein percentage, Milk fat yield, Milk protein yield, Milk caprylic acid content, Milk capric acid content, Milk myristic acid content, Milk myristoleic acid content, Milk pentadecylic acid content, Milk palmitic acid content, Milk palmitoleic acid content, Milk oleic acid content, Milk linoleic acid content, Milk linolenic acid content, Milk conjugated linoleic acid content, Milk C14 index, Milk C16 index, Milk C18 index, Curd yield, Curd solids yield, Curd firmness, Milk protein percentage, Milk alpha-lactalbumin percentage, Milk beta-lactoglobulin percentage, Milk alpha-S1-casein percentage, Milk casein percentage, Milk fat percentage, Milk saturated fatty acid content, Milk monounsaturated fatty acid content, Milk unsaturated fatty acid content, Milk polyunsaturated fatty acid content, Milk stearic acid content, Milk calcium content, Milk phosphorus content, Milk potassium content, Milk citrate content |
| 14 | Reproduction QTL | 1 | 0.5 - 0.5 | Conception rate |
| 16 | Health QTL | 1 | 46.2 - 46.2 | Clinical mastitis |
| 18 | Health QTL | 1 | 57 - 57 | Eosinophil number |
| 18 | Reproduction QTL | 17 | 13.3 - 57.1 | Stillbirth, Birth index, Calving ease, Calf size, Calving interval |
| 19 | Milk QTL | 8 | 36.7 - 36.7 | Milk caprylic acid content, Milk capric acid content, Milk lauric acid content, Milk myristic acid content, Milk palmitic acid content, Milk oleic acid content, Milk C14 index, Milk C18 index |
| 21 | Production QTL | 1 | 0 - 0 | Dry matter intake |
| 23 | Milk QTL | 1 | 26.9 - 26.9 | Milk conjugated linoleic acid content |
| 26 | Milk QTL | 11 | 2.9 - 2.9 | Milk caprylic acid content, Milk capric acid content, Milk lauric acid content, Milk myristic acid content, Milk myristoleic acid content, Milk palmitic acid content, Milk palmitoleic acid content, Milk stearic acid content, Milk C14 index, Milk C16 index, Milk C18 index |

### Cross-Trait Overlap and Functional Convergence

Cross-trait screening revealed varying degrees of functional overlap across biological levels (Figure 10). At the gene and protein levels, overlap was limited to a single entity (CCDC170 / coiled-coil domain containing 170) shared between MCE and MSB. In contrast, pathway and QTL levels displayed extensive cross-trait overlap.

**Figure 10.**
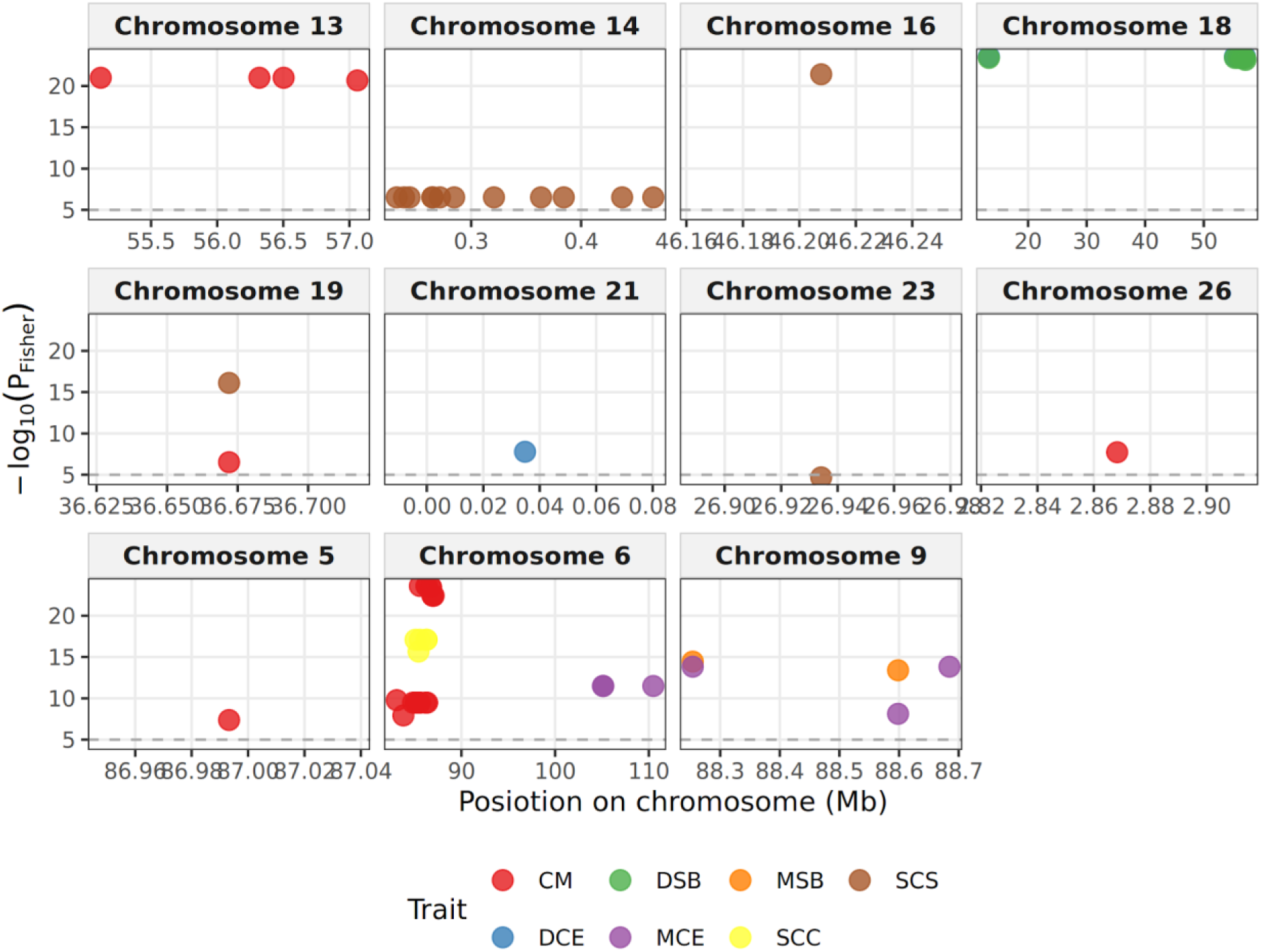
Chromosomal distribution of significant quantitative trait loci (QTL) identified across functional traits in dairy cattle.

**Figure 11.**
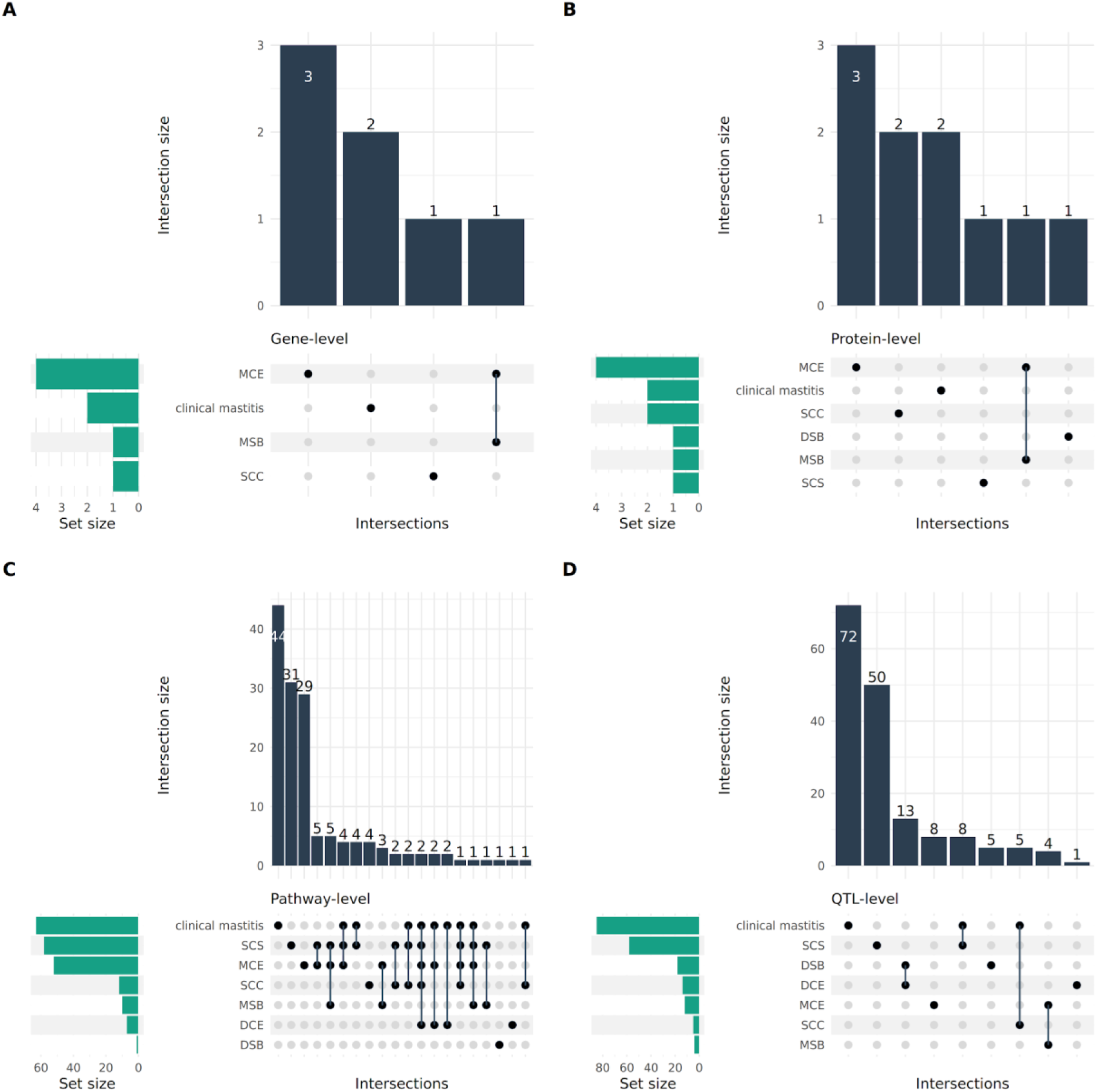
UpSet plots illustrating cross-trait functional overlap and shared biological entities across (A) genes, (B) proteins, (C) pathways, and (D) quantitative trait loci.

A total of 35 pathways showed cross-trait overlap (22 Reactome, 13 KEGG; Table 8). The most broadly shared pathways were Innate Immune System (R-BTA-168249) and Immune System (R-BTA-168256), both significant across 5 traits (CM, DCE, MCE, SCC, SCS). Signal Transduction (R-BTA-162582) and Metabolism (R-BTA-1430728) overlapped across 4 traits. At the QTL level, 30 QTL were shared across trait pairs (Table 9). Overlaps were observed between DCE and DSB, MCE and MSB, CM and SCC, and CM and SCS.

**Table 8.** Shared biological pathways across traits.

| ID | Description | Pathways |
| --- | --- | --- |
| R-BTA-168249 | Innate Immune System | CM, DCE, MCE, SCC, SCS |
| R-BTA-168256 | Immune System | CM, DCE, MCE, SCC, SCS |
| R-BTA-162582 | Signal Transduction | CM, MCE, MSB ,SCS |
| R-BTA-1430728 | Metabolism | CM, MCE, SCC, SCS |
| R-BTA-212436 | Generic Transcription Pathway | MCE, MSB, SCS |
| R-BTA-392499 | Metabolism of proteins | MCE, MSB, SCS |
| R-BTA-597592 | Post-translational protein modification | MCE, MSB, SCS |
| R-BTA-73857 | RNA Polymerase II Transcription | MCE, MSB, SCS |
| R-BTA-74160 | Gene expression (Transcription) | MCE, MSB, SCS |
| R-BTA-1280218 | Adaptive Immune System | CM, DCE, MCE |
| bta04020 | Calcium signaling pathway | CM, DCE ,MCE |
| R-BTA-112316 | Neuronal System | CM, MCE ,SCS |
| R-BTA-2262752 | Cellular responses to stress | CM , MCE , SCS |
| R-BTA-8953897 | Cellular responses to stimuli | CM, MCE, SCS |
| bta05207 | Chemical carcinogenesis - receptor activation | CM, MCE ,SCS |
| R-BTA-382551 | Transport of small molecules | CM, SCC ,SCS |
| R-BTA-425407 | SLC-mediated transmembrane transport | CM, SCC ,SCS |
| R-BTA-5688426 | Deubiquitination | MCE, MSB |
| bta04961 | Endocrine and other factor-regulated calcium reabsorption - | MCE, MSB |
| bta05200 | Pathways in cancer | MCE, MSB |
| R-BTA-194315 | Signaling by Rho GTPases | MCE, SCS |
| R-BTA-556833 | Metabolism of lipids | MCE, SCS |
| R-BTA-9006925 | Intracellular signaling by second messengers | MCE, SCS |
| R-BTA-9716542 | Signaling by Rho GTPases, Miro GTPases and RHOBTB3 | MCE, SCS |
| bta04081 | Hormone signaling | MCE, SCS |
| bta04080 | Neuroactive ligand-receptor interaction - | MSB, SCS |
| bta04972 | Pancreatic secretion | SCC, SCS |
| bta04976 | Bile secretion | SCC, SCS |
| bta04310 | Wnt signaling pathway | CM, DCE |
| bta05170 | Human immunodeficiency virus 1 infection | CM, DCE |
| R-BTA-425393 | Interactor (Protein) | CM, SCC |
| R-BTA-112315 | Transmission across Chemical Synapses | CM, SCS |
| bta00230 | Purine metabolism | CM, SCS |
| bta04714 | Thermogenesis | CM, SCS |
| bta04724 | Glutamatergic synapse | CM, SCS |

**Table 9.**
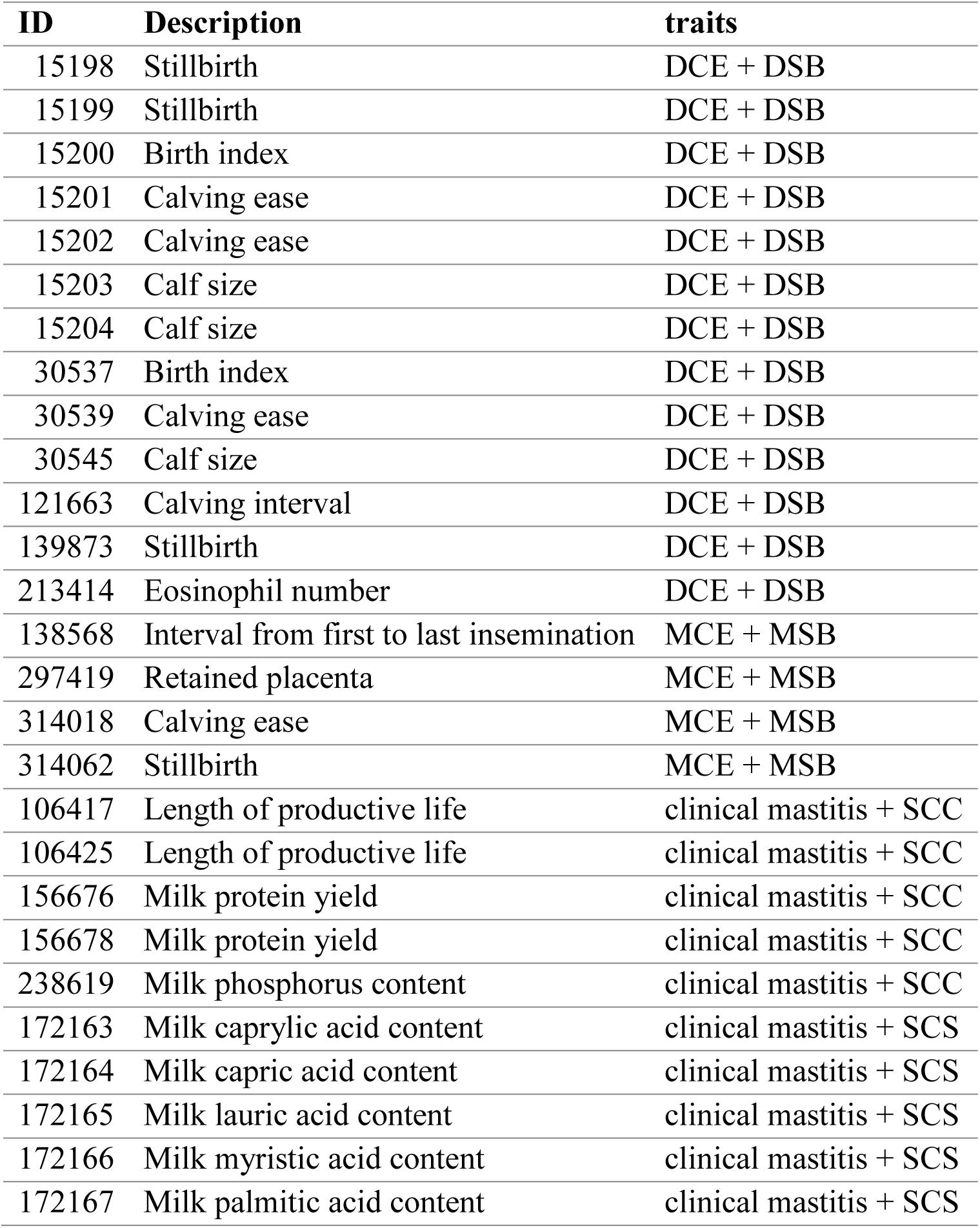
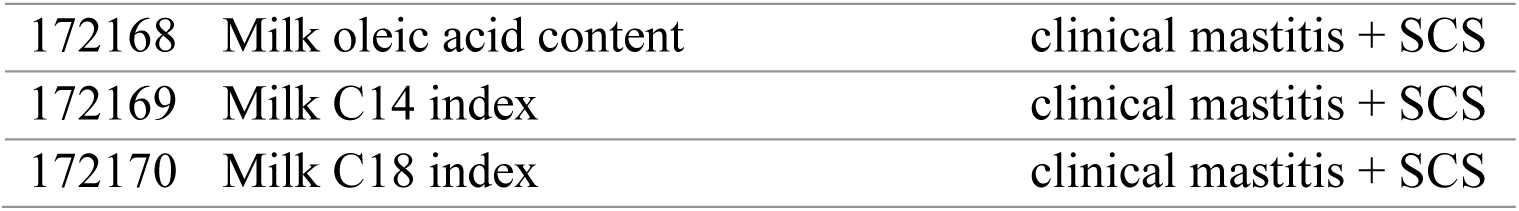
Shared quantitative trait loci (QTL) across traits.

## DISCUSSION

In this study, we combined a systematic review of published GWAS for calving performance and udder health traits in dairy cattle with a multi-level meta-analysis across 5 biological organizational tiers. Conventional SNP-level meta-analysis was uninformative due to minimal marker replication across studies: only 2 variants overlapped between studies, both originating from the same pair of publications, and neither achieved significance upon pooling. However, when variants were mapped to higher-order functional units-genes, proteins, pathways, and QTL-consistent association signals emerged for 6 of the 7 traits, and cross-trait overlap increased progressively with organizational level. These results support our hypothesis that GWAS results that rarely replicate at the individual variant level converge on common genes, biological pathways, and genomic regions.

The low SNP overlap reflects structural characteristics of the published literature rather than limitations of our dataset. Over a 16-year period, included studies utilized different genotyping platforms, reference genome assemblies, breeds, and sample sizes. Extensive linkage disequilibrium in cattle implies that the same underlying causal variant is frequently tagged by different markers across populations. Furthermore, published GWAS typically report only top associated markers rather than full summary statistics. Successful variant-level meta-analyses in cattle (Bouwman *et al*., 2018; Cai *et al*., 2024) relied on complete summary statistics, which are rarely available in literature-based meta-analysis settings.

### Biological Interpretation of Identified Signals

The strongest signal for MCE was ESR1, together with its genomic neighbor CCDC170, which was also significant for MSB. Both genes are located on BTA9 within the reproductive QTL hotspot identified in this study (88.3 to 88.7 Mb; calving ease, stillbirth, retained placenta). The prepartum surge in placental estrogens acting through estrogen receptors promotes myometrial contractility and cervical softening (Hoffmann and Schuler, 2002), establishing ESR1 as a major candidate gene for maternal delivery performance. Because the ESR1–CCDC170 genomic region is a recognized regulatory hotspot influencing ESR1 expression in humans (Veeraraghavan *et al*., 2014), CCDC170 variation may mark regulatory elements modulating ESR1 rather than representing a direct causal driver; expression QTL datasets such as the Cattle GTEx atlas (Liu *et al*., 2022) could clarify this relationship.

For udder health, DCK and TMPRSS11F (CM) and SLC4A4 (SCC) reside within a syntenic block containing GC, a primary candidate gene for the major mastitis QTL on BTA6 (Olsen *et al*., 2016; Cai *et al*., 2018). These genes map within the BTA6 hotspot (83.1 to 87.0 Mb) identified at the QTL level, indicating that they serve as positional proxies for this locus tagged by different markers across studies. This illustrates the utility of higher-level mapping where SNP-level integration fails. Similarly, the KRAB domain-containing protein associated with DSB and the concentration of BTA18 reproductive QTL for direct calving traits align with the established calving QTL on BTA18 (Dachs *et al*., 2023).

Pathway findings aligned with known trait biology. Mastitis-related traits were enriched for MAPK, PI3K-Akt, chemokine signaling, cell adhesion molecules, and neutrophil degranulation-pathways essential for neutrophil recruitment into milk during intramammary infection (Paape *et al*., 2003). Maternal calving ease was dominated by estrogen, prolactin, and calcium signaling, consistent with endocrine regulation of parturition and the role of periparturient hypocalcemia in dystocia (Goff, 2008). At the QTL level, the BTA14 hotspot for SCS reflects dense annotation of the DGAT1 region (Grisart *et al*., 2002) in Animal QTLdb, where milk production traits are heavily represented (Hu *et al*., 2022).

### Cross-Trait Overlap and Functional Convergence

Overlap at gene and QTL levels strictly adhered to the direct–maternal division of calving traits: CCDC170 and BTA9 QTL were shared between MCE and MSB, whereas BTA18 QTL were shared between DCE and DSB, consistent with documented genetic correlations between calving difficulty and stillbirth (Steinbock *et al*., 2003). Shared QTL between CM and SCC/SCS reflect high genetic correlations among udder health measures (Carlén *et al*., 2004). At the pathway level, immune and calcium signaling pathways were shared across calving and udder health traits. This convergence is biologically coherent, as periparturient immunosuppression and hypocalcemia link dystocia with heightened mastitis susceptibility (Mallard *et al*., 1998; Goff, 2008). However, broad Reactome pathways (e.g., Immune System, Signal Transduction) encompass numerous genes and may capture non-specific background signals; specific cascades such as calcium signaling or SLC-mediated transport provide higher functional resolution. Because cross-trait overlap at these levels may stem from distinct variants operating within common biological networks, we define these observations as functional convergence rather than direct genetic pleiotropy.

### Limitations and Implications

Fisher’s combination test assumes independent *P*-values that are uniformly distributed under the null hypothesis. Because published SNPs were pre-selected based on statistical significance and certain studies shared animal cohorts, combined *P*-values reflect cumulative evidence consistency across literature rather than unbiased global tests, and effect sizes remain susceptible to winner’s curse bias (Kraft, 2008). Selecting the minimum *P*-value per unit can favor larger genes, extensive pathways, and densely annotated genomic regions; linkage-disequilibrium–aware methods like MAGMA (de Leeuw *et al*., 2015) mitigate this bias but require full summary statistics. Furthermore, positional mapping may assign variants to neighboring non-causal genes, and annotation redundancies or retrotransposon-derived artifacts can inflate unit counts. Therefore, identified loci and pathways should be interpreted as prioritized candidate targets. Despite these limitations, multi-level functional mapping provides a practical framework for extracting reproducible biology from published GWAS literature. The identified genes, networks, and genomic hotspots can serve as biological priors in feature-selection genomic prediction models (MacLeod *et al*., 2016), and their shared immune and metabolic architecture supports joint selection strategies for reproductive and health traits in dairy cattle breeding programs.

## NOTES

### Supplemental Material

Supplemental material for this article is available at 10.5281/zenodo.23211744

### Ethical Approval

No human or animal subjects were used; therefore, ethical approval by an Institutional Animal Care and Use Committee or Institutional Review Board was not required.

## Nonstandard abbreviations

CE: calving ease
DCE: direct calving ease\
MCE: maternal calving ease
SB: stillbirth
DSB: direct stillbirth
MSB: maternal stillbirth
CM: clinical mastitis

